## Supplementary Information for "The online metacognitive control of decisions"

### Supplementary Methods 1: Optimal control policy for *Bayesian value denoising*

Under the oMCD framework, metacognitive control operates without knowing about the exact value computations that eventually modify the first- and second-order moments of the value representations. Rather, it relies on simplifying assumptions regarding the anticipated dynamics of $\mu(t)$ and $\sigma(t)$, which are tied to effort efficacy parameters (cf. Equations 3 and 4 in the main text). Here, we consider what the optimal stopping policy should be, would the control system know about the exact value computations that determine the dynamics of $\mu(t)$ and $\sigma(t)$.

Under the *Bayesian value denoising* scenario, the dynamics of $\sigma(t)$ actually conform to MCD's simplifying assumption (Equation 3). However, this does not hold for the dynamics of $\mu(t)$, because the impact of incoming noisy value signals on the value representations decreases with time. This eventually renders the transition probability density $p\left(\Delta\mu(t+1)\middle|\Delta\mu(t)\right)$ non stationary (cf. Equation 12 in the main text).

First, note that Equations 20-21 can be rewritten as instantaneous changes in the moments of the value representations, i.e.: $\mu(t+1)=\mu(t)+\breve{\delta}(t+1)$, where the instantaneous perturbation $\breve{\delta}(t+1)$ of the value mode is given by:

$$\breve{\delta}(t+1)=\frac{1}{\frac{\Sigma}{\sigma(t)}+1}\left(y(t+1)-\mu(t)\right) \quad \text{(A1)}$$

The total perturbation $\tilde{\delta}(t+1)$ in Equation 4 simply consists of the accumulated instantaneous perturbations in Equation A1, i.e.: $\tilde{\delta}(t+1)=\sum_{t'=1}^{t+1}\breve{\delta}(t')$. Importantly, the transition probability density $p\left(\Delta\mu(t+1)\middle|\Delta\mu(t)\right)$ is determined by the summary statistics of the perturbation $\breve{\delta}(t+1)$, conditional on $\mu(t)$. First, $E\left[\breve{\delta}(t+1)\middle|\mu(t)\right]=0$ because $E\left[y(t+1)\middle|\mu(t)\right]=\mu(t)$. Second, the variance of the instantaneous perturbation can be obtained from Equation A1:

$$E\left[\breve{\delta}(t+1)^2\middle|\mu(t)\right]=\frac{\Sigma+\sigma(t)}{\left(\frac{\Sigma}{\sigma(t)}+1\right)^2} \qquad \text{(A2)}$$

where $\sigma(t)$ can be written as a deterministic function of time (cf. Equation 20). Equation A2 states that the magnitude of instantaneous perturbations to value modes eventually become negligible as time increases, because $\sigma(t)$ tends towards 0. This implies that value signals that are sampled later in time are less likely to change one's decision, which is neglected by oMCD.

It follows that, under the *Bayesian value denoising* scenario, the exact transition probability density of the value mode difference is given by:

$$p\left(\Delta\mu(t+1)\middle|\Delta\mu(t)\right)=N\left(\Delta\mu(t),\frac{\Sigma+\sigma(t)}{\left(\frac{\Sigma}{\sigma(t)}+1\right)^2}\right) \quad \forall t\geq 1 \qquad \text{(A3)}$$

which is not stationary.

The ensuing "ideal" control policy then follows the exact same recursive backward induction steps than oMCD, having replaced oMCD's stationary transition probability density in Equation 12 in the main text with Equation A3.

So how different is the ideal control policy from oMCD's control policy?

Figure S1 below summarizes the results of a representative simulation series. (having set model parameters to $\alpha = 1$, $\nu = 2$, $\Sigma = 2$ and $\sigma_0 = 1$). We simulated 500 sample path trajectories of Bayesian value belief updates. Panel A shows the ensuing variance of instantaneous mode perturbations (Equation A2). For each belief trajectory, we extracted the invested resource, achieved confidence and net benefit under both oMCD and ideal policies.

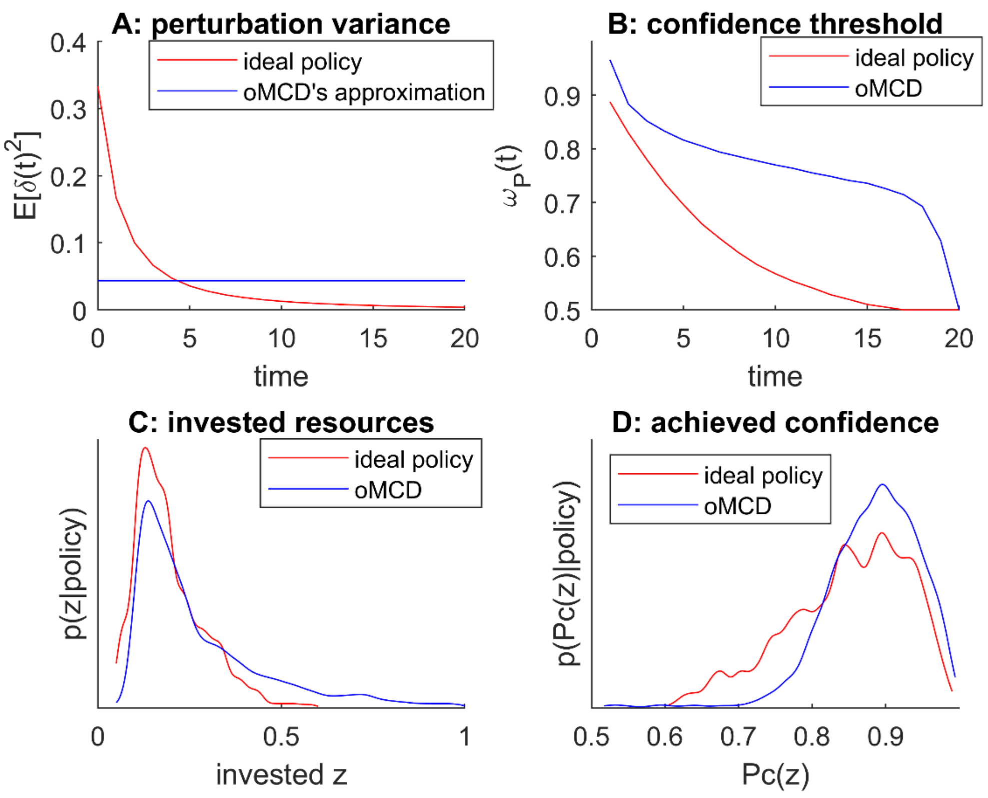


**Figure S1: Bayesian value denoising: exemplary comparison of oMCD and ideal control policies**. **A**: the variance of the instantaneous mode's perturbation $E\left[\breve{\delta}(t)^2\right]$ (y-axis, red: ideal policy, blue: oMCD approximation) is plotted against decision time $t$ (x-axis). **B**: the optimal confidence threshold $\omega_P^*(t)$ (y-axis) is plotted against decision time $t$ (x-axis). **C**: histogram of resource investments over 500 simulations of Bayesian value denoising trajectories, under both policies. **D**: histogram of achieved confidence over 500 simulations of Bayesian value denoising trajectories, under both policies.

Although both start at comparable levels, oMCD's confidence threshold dynamics lies above the ideal policy. This is essentially because oMCD underestimate the progressive dampening of the variance of the value modes' perturbation term. This increases the chance of a late bound hit, when compared to the ideal policy. This means that oMCD's control policy will induce stronger resource investments than the ideal policy (cf. panel C). However, oMCD decisions will achieve higher confidence than under the ideal policy (cf. panel D). We present a more exhaustive comparison of both policies in the Results section of the main text.

**Supplementary Methods 2: Optimal control policy for *progressive attribute integration***

As for *Bayesian value denoising*, the case of *progressive attribute integration* involves value computation details that oMCD neglects. In the latter case, the stochastic dynamics of the value mode $\mu(t)$ do comply with oMCD's static transition probability distribution (Equation 12). However, value variances $\sigma(t)$ do not evolve according to Equation 3. Predicting confidence under Equation 3 thus induces approximation errors that eventually deviate the optimal confidence threshold from the ideal control policy. So how does the ensuing ideal control policy look like?

Recall that the value variance dynamics obeys:

$$\sigma(t)=\sigma(t-1)-w_{K(t)}^{2}\varsigma_{K(t)} \qquad \text{(A4)}$$

where $K(t)$ is the index of the attribute that is sampled at time $t$ . Although value variances are necessarily decreasing over time, Equation A4 suggests that the dynamics of $\sigma(t)$ is stochastic. This is because $\sigma(t)$ is tied to the a priori unknown permutation order of attribute sampling $K(t)$. Now, since index permutations are equiprobable, the transition probability distribution of variances is stationary and writes:

$$p\left(\sigma(t)\middle|\sigma(t-1)\right)=\begin{cases}\dfrac{1}{k} & \text{if } \exists k':\sigma(t)=\sigma(t-1)-w_{k'}^{2}\varsigma_{k'}\\ 0 & \text{otherwise}\end{cases} \qquad \text{(A5)}$$

Equation A5 holds at any time, except for $t=T$ (time horizon), because then $\sigma(T)=0$ by construction (all the attributes have been sampled).

Although the transition probability distribution over value variances typically exhibits a sparse band-like structure, the induced value variance dynamics tends towards a dense distribution, even at low $k$. This is due to the cumulative effect of permutations over attribute sampling, which increases the number of admissible value variances in a quasi-exponential manner (at least over intermediate decision times).

Let us now summarize the ideal control policy.

To begin with, note that the net benefit $Q\left(a(t),\Delta\mu(t),\sigma(t)\right)$ is now formally a function of both moments of value representations. As we will see, this implies that the ideal policy cannot be written as a function of a single threshold on confidence. Nevertheless, this does not mean that oMCD's policy cannot be used as a quasi-optimal approximation to the ideal policy.

At $t=T$, $\sigma(T)=0$ thus confidence is maximal. Hence, the net benefit is: $Q\left(0,\Delta\mu(T),\sigma(T)\right)=R-\alpha\kappa T$, irrespective of $\Delta\mu(T)$.

At $t=T-1$, the ideal policy is $\pi^{*}(T-1)=0$ (stop) if $Q\left(0,\Delta\mu(T-1),\sigma(T-1)\right)>R-\alpha\kappa T$, and $\pi^{*}(T-1)=1$ (continue) otherwise ; and the optimal net benefit is:

$$Q^{*}\left(\Delta\mu(T-1),\sigma(T-1)\right)=\max\left\{Q\left(0,\Delta\mu(T-1),\sigma(T-1)\right),R-\alpha\kappa T\right\} \quad \text{(A6)}$$

For $t\leq T-2$, the ideal policy is:

$$\pi^{*}(t)=\begin{cases}0 \text{ if } Q\left(0,\Delta\mu(t),\sigma(t)\right)>E\left[Q^{*}\left(\Delta\mu(t+1),\sigma(t+1)\right)\middle|\Delta\mu(t),\sigma(t)\right]\\ 1 \text{ otherwise}\end{cases} \quad \text{(A7)}$$

and the optimal net benefit is:

$$Q^{*}\left(\Delta\mu(t),\sigma(t)\right)=\max\left\{Q\left(0,\Delta\mu(t),\sigma(t)\right),E\left[Q^{*}\left(\Delta\mu(t+1),\sigma(t+1)\right)\middle|\Delta\mu(t),\sigma(t)\right]\right\} \quad \text{(A8)}$$

where the conditional expectations in Equations A7 and A8 is taken under the transition probability distributions $p\left(\Delta\mu(t)\middle|\Delta\mu(t-1)\right)$ and $p\left(\sigma(t)\middle|\sigma(t-1)\right)$ over the value mode difference and the value variance, respectively.

The ideal policy for this MDP is obtained by iterating Equations A7 and A8 backward over time, i.e. from $t = T - 2$ to $t = 0$. Note that the ideal policy is to stop ($\pi^*(t) = 0$) whenever $\Delta\mu(t)$ is sufficiently large and/or $\sigma(t)$ is sufficiently small. How large $\Delta\mu(t)$ has to be to trigger the decision actually depends upon $\sigma(t)$, and reciprocally. This implies that the ideal policy frontier is in fact a unidimensional manifold on the 2D space spanned by the admissible ranges of $\Delta\mu(t)$ and $\sigma(t)$. A representative example of the ideal policy frontier can be eyeballed on panel A of Figure S2 (when setting: $\alpha = 1$, $v = 4$, $k = 8$ and sampling $\eta$, $\zeta$ and $w$ at random).

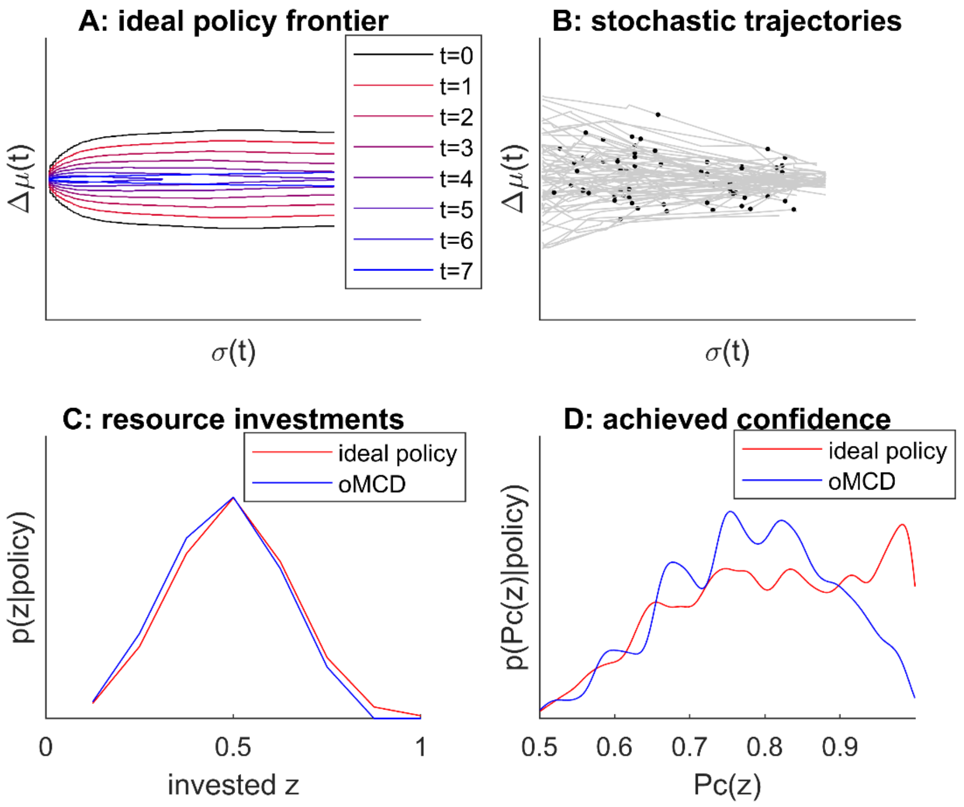


**Figure S2: Ideal policy for progressive attribute integration. A**: Ideal policy frontiers at different decision times (color code) are plotted on the 2D space spanned by the mode differences (x-axis) and value variances (y-axis). **B**: Exemplar sample paths of moments of value representations. Black dots show points at which the ideal policy triggers a decision. **C**: Histogram of invested resources over 1000 simulations of attribute integration trajectories, under both policies. **D**: Histogram of achieved confidence over 1000 simulations of attribute integration trajectories, under both policies.

The ideal policy is to continue as long as sample paths of moments of value representations lie within the ideal policy frontiers. In this example, the inner area of the critical region within which the ideal policy is to continue shrinks when time increases, which is qualitatively similar to oMCD's control policy (not shown).

Panels C and D of Figure S2 show the distributions of invested resources and achieved confidence for the same set of sample paths of value representation moments, when controlled using either the ideal control policy or oMCD's. One can see that both control policies tend to induce a very similar distribution of resource investment (cf. panel C). However, the ideal policy achieves higher confidence levels on average (cf. panel D). This means that, in this example, the average net benefit of oMCD will be lower than that of the ideal policy. An exhaustive comparison of both policies is presented in the Results section of the main text.

**Supplementary Methods 3: Optimal control policy with the aim of maximizing value**

The working assumption underlying MCD [1] is that decision confidence serves as the main benefit term of the resource allocation problem [2,3], where confidence is the subjective probability of having identified the best option. This apparently contrasts with standard treatments of value-based decision making, which equates the benefit of value-based decisions with the value of the chosen option [4–6]. Having said this, it is relatively easy to show that the two ideas are much more similar than one might intuitively think. This is the purpose of this Appendix.

To simplify the comparison, we assume that moments of value representations change according to Equations 2 and 3. In this context, oMCD provides the optimal control policy when the benefit of allocating resources is decision confidence. But what is the optimal control policy rule when setting the benefit to the expected value of the chosen option? In what follows, we refer to this policy as *max(value)*.

Recall that the chosen option is the option that has the maximum expected value. Now, the maximum $M = \max\{X_1, X_2\}$ of two arbitrary variables $X_1$ and $X_2$ can be written as a function of the absolute difference between the two, i.e.: $M = \frac{1}{2}(X_1 + X_2) + \frac{1}{2}|X_1 - X_2|$. This implies that the value of the chosen option is simply $\max_i\{\mu_i(\tau)\} = \frac{1}{2}\bar{\mu}(\tau) + \frac{1}{2}|\Delta\mu(\tau)|$, where $\tau$ is the decision time and $\bar{\mu}(\tau) = \frac{1}{2}(\mu_1(\tau) + \mu_2(\tau))$ is the mean expected value (across options).

Thus, the net benefit that the decision system would obtain at time $t$ is given by:

$$Q(a(t), \Delta\mu(t), \bar{\mu}(t)) = \begin{cases} \bar{\mu}(t) + \dfrac{1}{2}|\Delta\mu(t)| - C(t) & \text{if } a(t) = 0 \\ 0 & \text{otherwise} \end{cases} \quad \text{(A10)}$$

where $C(t)$ is the cost of decision time (cf. Equation 7), and we have removed the dependency to time for notational convenience.

At $t = T$, the system stops deliberating; it then receives a net benefit $Q(0, \Delta\mu(T), \bar{\mu}(T))$.

At $t = T - 1$, *max(value)*'s optimal policy is:

$$\pi^*(T-1) = \begin{cases} 0 \text{ if } \Delta Q(T-1) > 0 \\ 1 \text{ otherwise} \end{cases} \tag{A11}$$

where the control criterion $\Delta Q(T-1)$ is given by:

$$\begin{aligned} \Delta Q(T-1) &= \underbrace{Q(0, \Delta\mu(T-1), \bar{\mu}(T-1))}_{\text{net benefit of stopping}} - \underbrace{E\left[Q(0, \Delta\mu(T), \bar{\mu}(T)) \middle| \mu(T-1)\right]}_{\text{expected net benefit of waiting}} \\ &= \frac{1}{2}\left(\left|\Delta\mu(T-1)\right| - E\left[\left|\Delta\mu(T)\right| \middle| \mu(T-1)\right]\right) - \left(C(T-1) - C(T)\right) \end{aligned} \tag{A12}$$

The conditional expectation in the RHS of Equation A12 is taken under the transition law of value modes (cf. Equation 12 in the main text).

Importantly, Equation A12 states that the control criterion of *max(value)* is independent of the mean expected value $\bar{\mu}$. This is because $E\left[\bar{\mu}(T) \middle| \mu(T-1)\right] = \bar{\mu}(T-1)$, and hence the difference vanishes in $\Delta Q(T-1)$. In turn, the control criterion of *max(value)* is only a function of the absolute value mode difference $|\Delta\mu|$, which is the main driver of decision confidence. In other words, deriving *max(value)*'s optimal control policy reduces to deriving an optimal threshold $\omega_\Delta(T-1)$ on $|\Delta\mu(T-1)|$, i.e.:

$$\pi^*(T-1) = \begin{cases} 0 \text{ if } \left|\Delta\mu(T-1)\right| > \omega_\Delta(T-1) \\ 1 \text{ otherwise} \end{cases} \tag{A13}$$

where the optimal threshold $\omega_\Delta(T-1)$ is:

$$\omega_{\Delta}(T-1) = E\left[\left|\Delta\mu(T)\right|\left|\mu(T-1)\right.\right] + 2\left(C(T-1) - C(T)\right) \quad \text{(A14)}$$

This is very similar to oMCD's control rule, which derives from comparing decision confidence at $t = T - 1$ to the expected decision confidence at $t = T - 1$, both of which being monotonic functions (more precisely: sigmoid mappings) of $\left|\Delta\mu(T-1)\right|$. The similarity between the net benefit of control and waiting under both frameworks is exemplified in Figure S3 below, where we have set $\alpha = 1$, $\beta = 1$, $\gamma = 1$, $\nu = 1$, $\kappa = 0.05$ and $\sigma(T-1) = 0.1$.

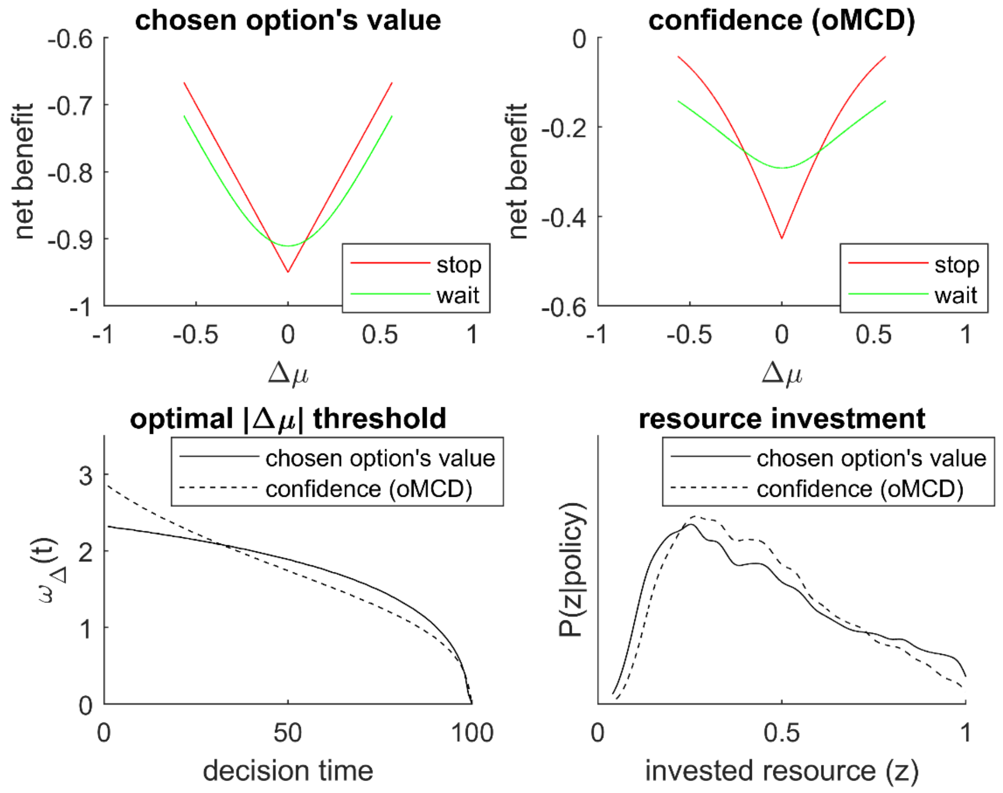


**Figure S3. Comparison of optimal control policies.** The net benefit of control (y-axis, red) and waiting (y-axis, green) are plotted as a function of the current difference in value means $\Delta\mu$ at $t = T - 1$ (x-axis). **A**: the benefit of decisions is set as the value of the chosen option. **B**: the benefit of decisions is set as confidence (oMCD's policy).

Although the exact value mode difference that triggers a decision is not the same under both settings of decision benefits, the net benefit functions that determine the optimal control policy behave qualitatively similarly. So are the dynamics of optimal control thresholds similar under both frameworks? At this point, we note that oMCD's net benefit function is sensitive to the variance of value representations, which is not the case for *max(value)*.

From an algorithmic perspective, the full derivation of *max(value)*'s optimal threshold is very similar to that of oMCD:

- At $t = T$, $\omega_\Delta(T) = 0$ by convention.
- At $t = T - 1$, the optimal threshold $\omega_\Delta(T-1)$ is given by Equation A14.
- At $t < T - 1$, the optimal net benefit is derived under Bellman's optimality principle:

$$\begin{aligned} Q^*(\Delta\mu(t)) &= Q(\pi^*(t), \Delta\mu(t), \bar{\mu}(t)) \\ &= \max\left\{Q(0, \Delta\mu(t), \bar{\mu}(t)), E\left[Q^*(\Delta\mu(t+1)) \middle| \Delta\mu(t)\right]\right\} \end{aligned} \quad \text{(A18)}$$

  where the expectation is taken under the transition density of value modes (cf. Equation 12). Then the optimal threshold $\omega_\Delta(t-1)$ is the absolute value mode difference $\Delta\mu^*(t-1)$ such that: $Q(0, \Delta\mu^*(t), \bar{\mu}(t)) = E\left[Q^*(\Delta\mu(t)) \middle| \Delta\mu^*(t-1)\right]$. This defines a recurrence relationship that can be applied backward in time.

Importantly, the ensuing threshold dynamics depends upon cost parameters ($\alpha$ and $\nu$), as well as effort intensity $\kappa$ and type #2 effort efficacy ($\gamma$). However, in contrast to oMCD, it will be insensitive to type #1 effort efficacy ($\beta$). We provide an exhaustive comparison of both policies in the Results section of the main text.
